# Reduced S-acylation of SQSTM1/p62 in Huntington disease is associated with impaired autophagy

**DOI:** 10.1101/2023.10.11.561600

**Authors:** F. Abrar, M.C. Davies, A. Kumar, A. Dang, Y.T.N. Nguyen, J. Collins, N. Caron, J.S. Choudhary, S.S. Sanders, M.O. Collins, M.R. Hayden, D.D.O. Martin

## Abstract

Disruption of macroautophagy/autophagy has emerged as a common feature in many neurodegenerative diseases. Autophagy is a membrane-dependent pathway that requires many key regulators to quickly localize on and off membranes during induction promoting membrane fusion. Previously, our bioinformatic approaches have shown that autophagy and Huntington disease (HD) are enriched in S-acylated proteins. S-acylation involves the reversible addition of long chain fatty acids to promote membrane binding. Herein, we show that inhibition of S-acylation regulates the abundance of several key regulators of autophagy and leads to a partial block of autophagic flux. We show that the autophagy receptor SQSTM1/p62 (sequestosome 1) is S-acylated and directed to the lysosome. Importantly, we see that SQSTM1 S-acylation is significantly reduced in HD patient and mouse model brains, thus providing a novel mechanism for the generation of empty autophagosomes previously seen in HD models and patient cells.

## Introduction

Protein mislocalization and proteostasis deficiency are hallmarks of neurodegeneration^1–3^. The cell has two primary ways to eliminate misfolded and aggregating proteins; the unfolded protein response and autophagy. The unfolded protein response requires the proteasome but becomes quickly overwhelmed in diseases such as Huntington Disease (HD), amyotrophic lateral sclerosis (ALS), and other neurodegenerative diseases^2,4–6^; autophagy then becomes the bulk protein clearance pathway. Autophagy generally refers to a pathway that allows the delivery of damaged or superfluous organelles and toxic proteins to the lysosome for degradation and recycling^7^. Autophagy can be further classified into three broad categories; macroautophagy (hereafter referred to as autophagy), chaperone-mediated autophagy (CMA) and microautophagy^7^. A common theme among these pathways is that they involve dynamic changes in membranes and/or protein interactions with those membranes. However, in many cases, how autophagy regulators quickly transition from the cytosol to membranes is generally unknown or overlooked.

Previously, we provided the first unbiased bioinformatics analysis showing that S-acylation of proteins is enriched in neurodegenerative disease and proteostasis deficiencies including HD and ALS^8^. S-acylation refers to the reversible and covalent addition of saturated fatty acids to cysteine residues through a thioester bond^8,9^. The most common form involves palmitate and is generally referred to as S-palmitoylation, or simply palmitoylation. The hydrophobic moiety is primarily involved in membrane binding, but can also promote protein-protein interactions, conformational changes, or protein-folding^10,11^. Much like phosphorylation, S-acylation is reversible and dynamic and is regulated by diverse set of enzymes that add or remove palmitate from proteins; palmitoyl acyltransferases (PATs) and acyl protein thioesterases (APTs), respectively^12^. Thus, S-acylation provides an ideal mechanism for quickly redirecting proteins on and off membranes. When S-acylation is disrupted, it leads to protein mislocalization, decreased protein turnover and aggregation^5^.

Previous work has implicated a potential role for palmitoylation in autophagy^13,14^. Palmitate treatment increases LC3-I to LC3-II conversion as well as decreases SQSTM1 levels, all signs that autophagy levels are increased^14^. The use of the tandem LC3 reporter with palmitate treatment, shows a decrease in the number of autophagosomes and an increase in autolysosomes^14^; again indicating an overall increase in autophagy, and highlighting a link between palmitoylation and autophagy. In addition, we recently showed that the majority of autophagy regulatory genes encode a proteo form that has been detected by low- or high-throughput methods^5^, by comparing the S-acylation database SwissPalm^15^ and the Autophagy Regulatory Network^16^.

Autophagy induction requires the recruitment of MCOLN3/TRPML3, a Ca^2+^-permeable cation channel, to provide Ca^2+^ for autophagosome biogenesis^13^. It was recently shown that this channel requires palmitoylation at the C terminus, a modification essential for the trafficking and function of the channel in autophagy. When HeLa cells are starved of nutrients, MCOLN3/TRPML3 is activated alongside palmitoylation, with palmitoylation disruption resulting in activation of MCOLN3/TRPML3 and autophagy being abolished^13^. Recently, we identified multiple components of the MTOR complex 1 that are palmitoylated in human embryonic kidney 293T (HEK293T) cells^17^. Again, this work implicates a direct role of palmitoylation in autophagy. However, further investigation of the exact role palmitoylation plays in the regulation of autophagy is needed.

Herein, we show that S-acylation may provide a mechanism for autophagy regulators to quickly and reversibly relocalize on and off membranes. Furthermore, we find that the autophagy receptor SQSTM1/p62 is S-acylated and directed to the lysosome. Finally, we see that SQSTM1 S-acylation is significantly reduced in HD patient and mouse model brains. This finding may provide a mechanism for the production of empty autophagosomes previously seen in HD models and patient cells^18,19^.

## Materials and Methods

### Materials

This study examined de-identified archived brain tissue samples from the Huntington Disease BioBank at the University of British Columbia. All samples were collected, stored, and accessed with informed consent and approval of The University of British Columbia/Children’s and Women’s Health Centre of British Columbia Research Ethics Board (UBC C&W REB H06-70467 and H06-70410). Tissues used were from patients of similar age at death, CAG size, and post-mortem intervals. Male and female samples were included in controls and diseased samples. HA-tagged SQSTM1 and RFP-LC3 were gifts from Qing Zhong (Addgene, 28027)^20^ and Tamotsu Yoshimori (Addgene, 21075)^21^, respectively. All point mutations for SQSTM1-HA were generated by site-directed mutagenesis by TopGene Biotech (Ontario, Canada). Cathepsin B-BFP (CTSB-BFP) was a kind gift from the Davidson laboratory stocks. Unless otherwise indicated, all antibodies were acquired from Cell Signaling Technology or Millipore-Sigma.

### Mice

All mouse experiments were carried out in accordance with protocols approved by the UBC Committee on Animal Care and the Canadian Council on Animal Care (Animal protocol A07-0106). Mice were derived from in-house breeding pairs, maintained under a 12 h light:12 h dark cycle in a clean facility and given free access to food and water except where indicated (for fasting). As previously described, mice were fasted for 24 h with full access to water to induce autophagy in the brains^22^. YAC128 mice were on an FVB/N background and mixed sexes were analyzed (no sex differences were observed).

### Statistics

Unless otherwise indicated, statistical significance was assessed using Student’s t-test for comparison of two groups, one-way ANOVA with post-hoc Tukey’s correction for the comparison of one variable between more than two groups, and two-way ANOVA with post-hoc Bonferroni correction for the comparison of two variables between groups. Variances between groups were similar. All analyses were performed using the GraphPad Prism 5.01 or higher software package. Graphs are presented as standard error of the mean (SEM).

### Cell culture and SILAC labelling

HeLa cells were grown in normal DMEM (Gibco) supplemented with 1% penicillin-streptomycin (Gibco). Where indicated, induction of autophagy in HeLa cells was conducted by application of Earle’s Balanced Salt Solution (EBSS; Thermo Fisher, 24010043). Inhibition of autophagy was conducted by application of 500 nM bafilomycin A_1_ (BafA1; Alfa Aesar, J61835).

For SILAC experiments, DMEM lacking lysine and arginine (Dundee Cell Products) was supplemented with 40 mg/L lysine and 84 mg/L arginine of normal, light isotopic composition (Sigma) for the K0/R0 population, and heavy isotopes (4, 4, 5, 5-D4 L-lysine and 13C6 L-arginine, Cambridge Isotope Laboratories (CIL)) for the K4/R6 population. The medium was supplemented with 10% FBS (Gibco) that had been dialyzed at 1 kDa (SpectraPor) overnight in 150 mM NaCl to remove traces of normal weight isotopic amino acids. HeLa cells treated with either DMSO (Light, K0/R0) or 50 µM palmostatin B (heavy K4/R6) for 18 h.

### Quantitative proteome profiling of drug-treated cells

Cell pellets from each condition were lysed with a 4% SDS-containing extraction buffer, as previously described^23^, and pooled and 3 biological replicates were prepared and analyzed for each treatment group. Thirty µg of each pooled sample was separated on a 12% SDS-PAGE gel to fractionate proteins for quantitative proteome profiling. Protein gels were stained overnight with colloidal Coomassie Brilliant Blue (Sigma). Each lane was excised into 9 sections that were destained and in-gel digested overnight using trypsin (sequencing grade; Roche). Peptides were extracted from gel bands twice with 50% acetonitrile:0.5% formic acid and dried in a SpeedVac (Thermo). Peptides were resuspended in 0.5% formic acid and were analyzed by LC-MS/MS analyses using an Ultimate 3000 RSLC Nano LC System (Dionex) coupled to an LTQ Orbitrap Velos hybrid mass spectrometer (Thermo Scientific) equipped with an Easy-Spray (Thermo Scientific) ion source. Peptides were desalted on-line using a capillary trap column (Acclaim Pepmap100, 100 μm, 75 μm × 2 cm, C18, 5 μm (Thermo Scientific)) and then separated using a 180 min RP gradient (4–30% acetonitrile/0.1% formic acid) on an Acclaim PepMap100 RSLC C18 analytical column (2 μm, 75 μm id x 50 cm, [Thermo Scientific]) with a flow rate of 0.3 μl/min. The mass spectrometer was operated in standard data dependent acquisition mode controlled by Xcalibur 2.2. The instrument was operated with a cycle of one MS (in the Orbitrap) acquired at a resolution of 60,000 at m/z 400, with the top 15 most abundant multiply-charged (2 + and higher) ions in a given chromatographic window were subjected to CID fragmentation in the linear ion trap. An FTMS target value of 1e6 and an ion trap MSn target value of 10000 were used. Dynamic exclusion was enabled with a repeat duration of 30 s with an exclusion list of 500 and exclusion duration of 60 s. Lock mass of 401.922 was enabled for all experiments.

### Mass spectrometry data analysis

Data were analyzed using MaxQuant version 1.4.1.2. MaxQuant processed data were searched against a UniProt (downloaded October 2014) human sequence database using the following search parameters: trypsin with a maximum of 2 missed cleavages, 7 ppm for MS mass tolerance, 0.5 Da for MS/MS mass tolerance, with acetyl (Protein N-term) and oxidation (M) as variable modifications and carbamidomethyl (C) as a fixed modification. Light (K0/R0) and heavy (K4/R6) SILAC labels were specified for SILAC quantification. A protein FDR of 0.01 and a peptide FDR of 0.01 were used for identification level cut offs. Match between runs with a 2-min retention time window was enabled. Data were filtered so that at least 3 valid normalized SILAC ratios were present and two sample t-testing was performed with a permutation-based FDR calculation in Perseus (1.4.1.3).

### S-acylation assays

S-acylation using the acyl-resin assisted capture (Acyl-RAC) and immunoprecipitation acyl-biotin exchange (IP-ABE) assays was detected as described previously^24–26^. Briefly, cells were lysed in the presence of 100 mM NEM and incubated at 4°C overnight to alkylate free thiols. Following chloroform-methanol precipitation, 150 µg of lysate was used to react with the thiol reactive acyl-resin assisted capture beads (Thiopropyl Sepharose 6B, T8387) in the presence of hydroxylamine (HAM), which facilitates the exchange of thioester linked fatty acids with beads. Detection of SQSTM1 S-acylation was achieved using the mouse anti-SQSTM1/p62 (MBL, M162-3) antibody. For IP-ABE of endogenous SQSTM1, the protein was immunoprecipitated using the same antibody and detected with rabbit anti-SQSTM1 (Enzo Life Sciences, BML-PW9860). For HA-tagged proteins, SQSTM1-HA was immunoprecipitated with rat anti-HA (3F10) and detected with mouse anti-HA (12CA5). S-acylation was detected using Alexa Fluor 680-conjugated streptavidin (Invitrogen, S-32358).

### Microscopy

#### Endogenous SQSTM1 and tandem LC3 reporter

Cells were cultured in 12-well plates (Greiner Bio-One - 665180) containing coverslips (VWR - 631-0152) until ∼70% confluence. Media was removed and cells were washed with PBS three times, before cells were fixed with 4% PFA (Sigma-Aldrich - 158127) or ice-cold 100% methanol (VWR - BDH1135-4LG) for 20 min at room temperature (RT). Washing in PBS was conducted before permeabilization with PBS +0.3% Triton X-100 for 15 min or ice-cold 100% methanol. Cells were washed before blocking in PBS +0.01% Triton X-100 and 0.2% fish skin gelatin (Sigma-Aldrich - G7765) for 1 h at RT. Blocking buffer was removed and primary antibody was applied in blocking solution for 1 h at RT or 4°C overnight. Cells were washed in PBS three times then fluorescent secondary antibody was applied in blocking solution for 2 h in the dark. Cells were washed in PBS three times before a final wash in dH_2_0. Coverslips were removed from wells and air-dried before inversion onto DAPI-Fluoromount-G (Southern Biotech, 0100) on microscopy slides (J. Melvin Freed Brand, 7525M). Imaging was performed at the Wolfson Light Microscopy Facility at The University of Sheffield, using a NIKON A1 Confocal Microscope. The following wavelengths were used 405 nm (blue), 488 nm (green), 561 nm (red) and 595 nm (cyan). Images were acquired using a 60x objective lens and the NIS elements program (Nikon instruments) in an ND2 format. For all samples, no primary and no secondary controls were used to threshold the exposure at each individual laser wavelength. Puncta were counted in FIJI using the Cell Counter (Version - ImageJwin-64)^27^. Results were analyzed on GraphPad Prism 7. Statistical significance was accepted at p<0.05. Results are reported as means ± standard error of the mean.

#### Localization of SQSTM1-HA

HeLa cells were plated on poly-D-lysine-coated glass coverslips and transfected with SQSTM1-HA, RFP-LC3, and CTSB-BFP using XtremeGene (Roche). Rapamycin (100 nM) was added to induce autophagy and 400 nM BafA1 was used to inhibit the fusion of autophagosomes and lysosomes to prevent degradation of SQSTM1-HA. Cells were fixed in 4% paraformaldehyde. Rat anti-HA and anti-rat Alexa Fluor 488 were used to detect SQSTM1-HA and imaged on an SP8 Leica microsystems confocal microscope.

## Results

### Inhibition of depalmitoylation disrupts autophagy

To assess the extent of regulation of the autophagy network by S-acylation we measured changes in the proteome in response to global inhibition of depalmitoylation enzymes. Light and heavy SILAC labelled HeLa cells were treated with 0.1 % DMSO as a control or 100 µM palmostatin B for 18 h. Quantitative mass spectrometry of tryptic digests from replicate sets of control and palmostatin B samples revealed that 273 proteins were increased in abundance after palmostatin B treatment (Supplementary Table 1, Figure 1). Out of the top 15 upregulated proteins with highest fold change, six were involved in the autophagic process including SQSTM1 and LC3B.

**Figure 1.**
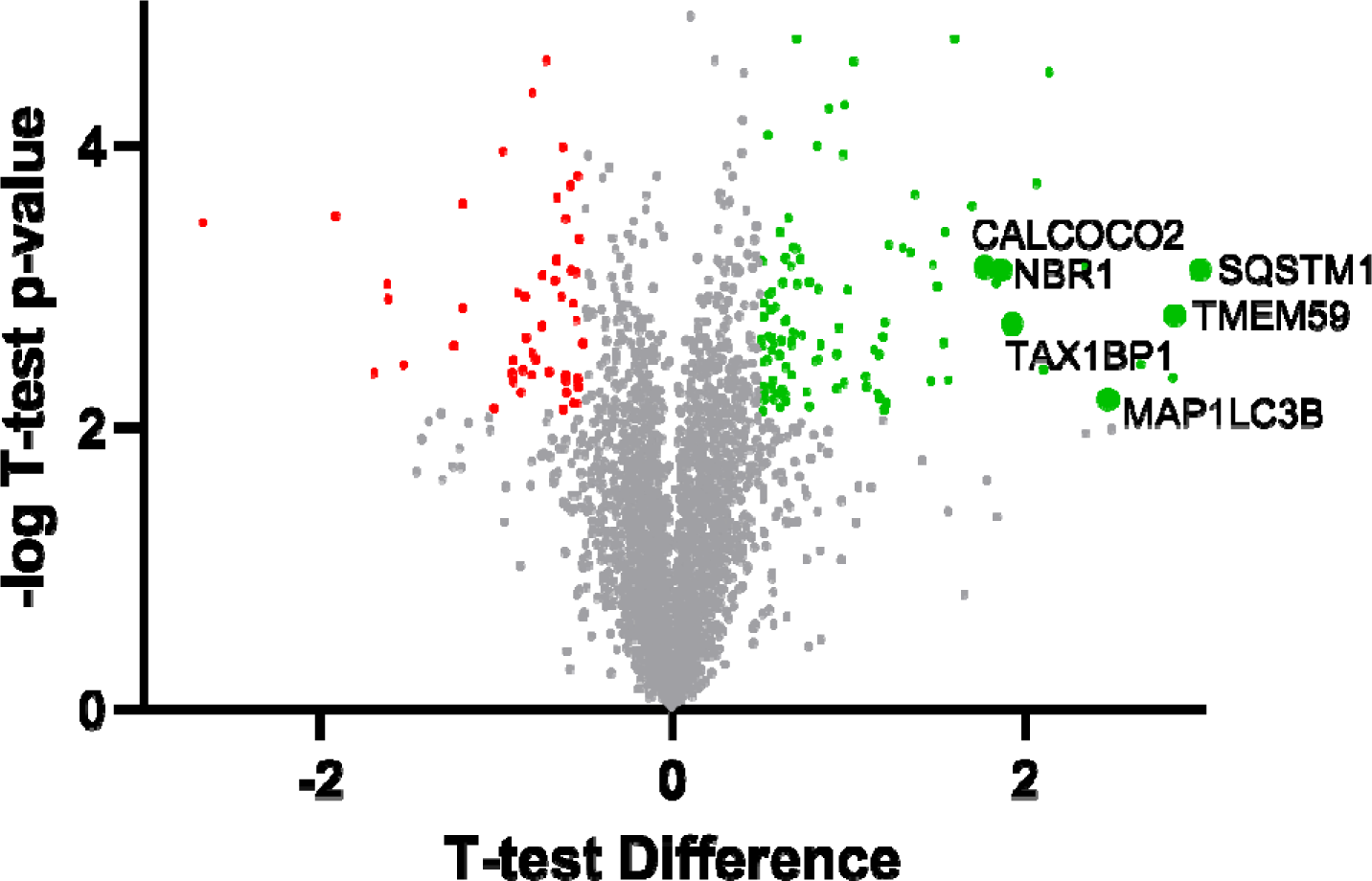
Palmostatin B treatment upregulates components of the autophagy pathway. Volcano plot of protein abundance changes in response to treatment of HeLa cells with 50 mM palmostatin B for 18 h. Control and palmostatin B-treated cells were SILAC labelled with light (K0/R0) and heavy (K4/R6) isotope labelled lysine and arginine to enable precise quantification of protein abundance changes. Significantly regulated proteins (red and green) were determined using two-way testing with a permutation-based FDR of 0.05 and a Log2 t-test difference filter of 0.5.

We next assessed the effect of palmostatin B treatment on autophagy induction (4 h EBSS), with and without blockage of late-stage autophagy using 500 nM BafA1. To measure autophagy, levels of SQSTM1 and LC3-II were assessed via immunoblotting and immunofluorescence microscopy. In the absence of palmostatin B, EBSS treatment increased LC3-II and SQSTM1 levels after as expected (Figure 2). However, palmostatin B treatment led to a reduction in SQSTM1 and LC3-II levels in EBSS versus control conditions and the addition of BafA1 to block turnover, increased the level of LC3-II. Together these data indicate that inhibition of palmitoylation causes a partial block in autophagy.

**Figure 2.**
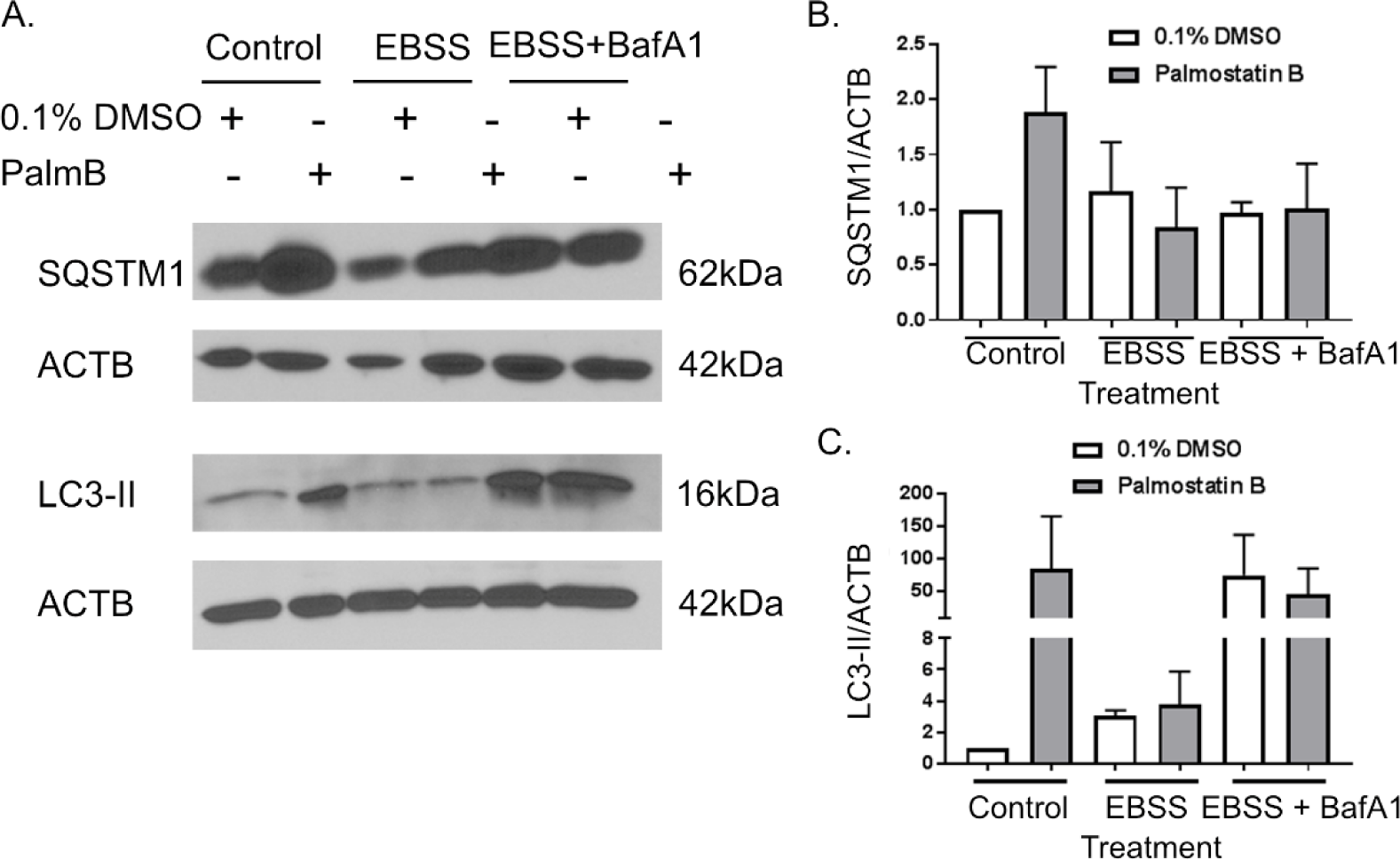
Palmostatin B alters autophagy in HeLa cells, assessed through LC3B and SQSTM1 immunoblotting. HeLa cells were treated with 0.1% DMSO or 100 μM palmostatin B (Palm. B) for 18 h. Autophagy was induced by 4-h EBSS treatment or induced and inhibited by 4-h EBSS/2-h treatment with 500 nM BafA1. A. Representative images of immunoblotting conducted for autophagy markers SQSTM1 and LC3-II with and without Palm. B treatment and autophagy induction. B. Quantification of immunoblots for SQSTM1 normalized to ACTB/β-actin. Quantification was conducted using ImageJ software (mean + SEM from three independent experiments; Mann Whitney test). C. Quantification of immunoblots for LC3-II normalized to ACTB. Quantification was conducted using ImageJ software (mean + SEM from three independent experiments; Mann Whitney test).

While the levels of LC3-II between different samples is a good measure of autophagic flux differences, we sought to determine if palmostatin B treatment changes the number of autophagosomes by measuring the number of LC3 puncta using immunofluorescence microscopy. Palmostatin B treatment did not affect SQSTM1 puncta (Figure 3) but it did increase the number of LC3B puncta per cell (Figure 4). To assess the exact change in autophagy, a tandem LC3 reporter was employed. There were no changes in autophagosome levels in any of the conditions for control versus palmostatin B-treated samples. However, there was an increase in the number of autolysosomes after palmostatin B treatment (Figure 5). This increase in the latter marker of autophagy suggests a late-stage blockage in autophagy which is in line with our immunoblotting data.

**Figure 3.**
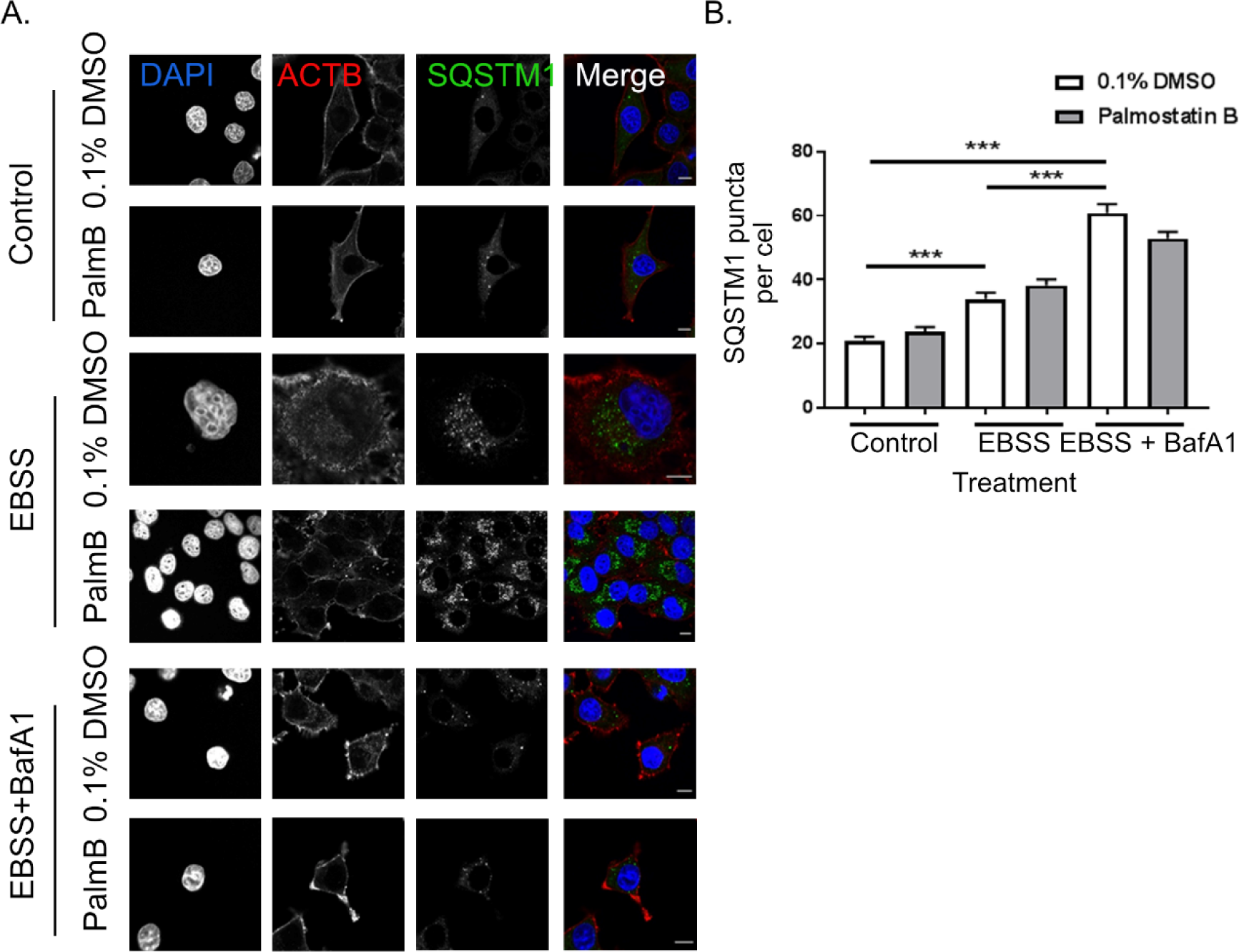
HeLa cells were treated with 0.1% DMSO or 100 μM palmostatin B (Palm. B) for 18 h. Autophagy was induced by 4-h EBSS treatment or induced and inhibited by 4-h EBSS and 2-h 500nM BafA1. **A.** Representative images of immunofluorescence conducted for SQSTM1 and ACTB after Palm. B and autophagy treatment. **B.** Quantification of the number of SQSTM1 puncta per cell. Quantification was conducted on ImageJ software (mean + SEM from three independent experiments; Mann Whitney test; N (cells) Control: 154, Control + Palm. B: 184, EBSS: 169, EBSS + Palm. B: 103, EBSS+BafA1: 125, EBSS+BafA1 + Palm. B: 120) *** p<0.001. Scale bars: 10 µm.

**Figure 4.**
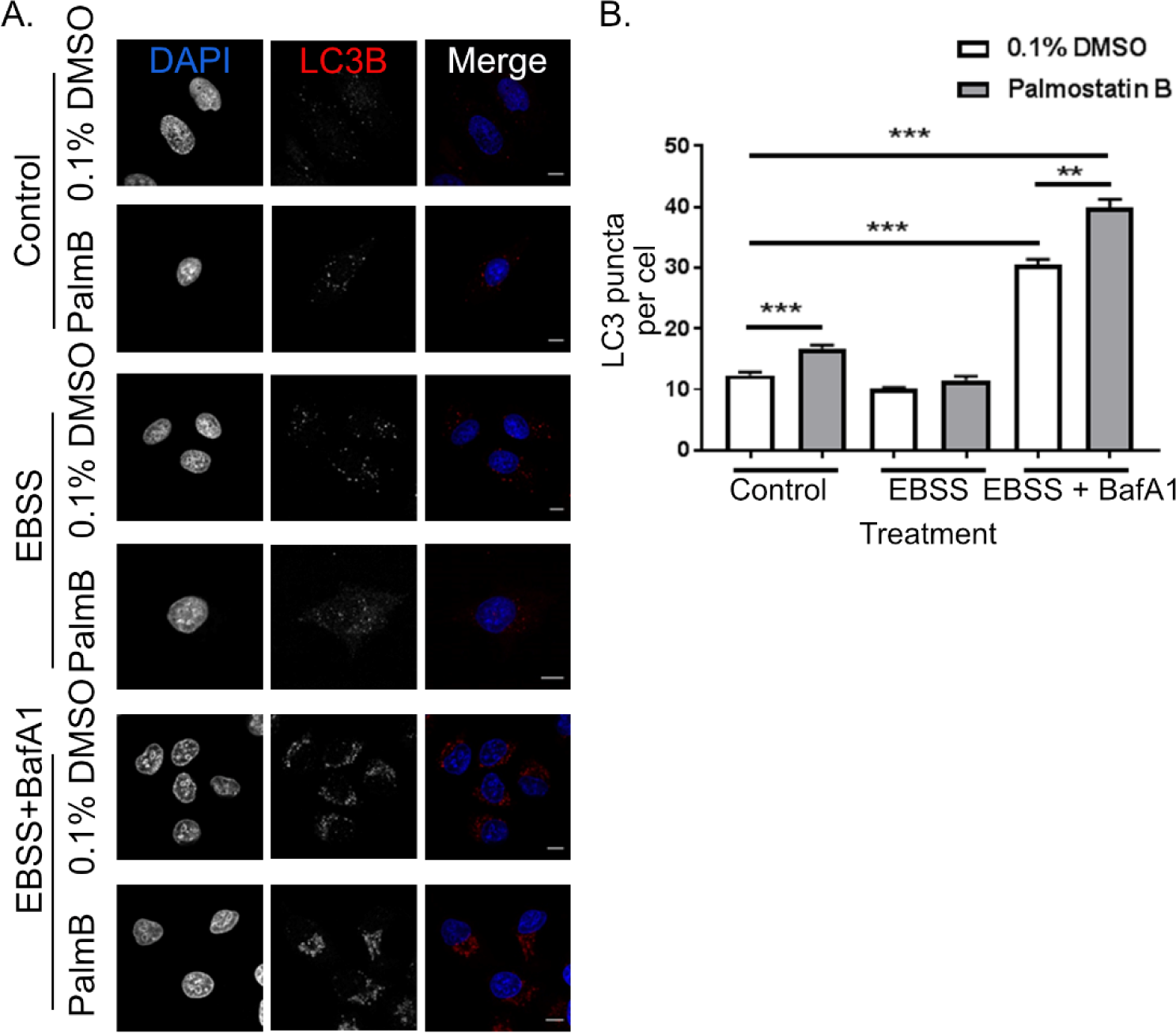
Palmostatin B alters autophagy in HeLa cells, assessed through LC3B immunofluorescence microscopy. HeLa cells were treated with 0.1% DMSO or 100 μM palmostatin B (Palm. B) for 18 h. Autophagy was induced by 4h EBSS treatment or induced and inhibited by 4h EBSS and 2h 500nM BafA1. **A.** Representative images of immunofluorescence conducted for LC3B after Palm. B and autophagy treatment. **B.** Quantification of LC3B puncta per cell. Quantification was conducted using ImageJ software (mean + SEM from three independent experiments; Mann Whitney test; N (cells) Control: 167, Control + Palm. B: 155, EBSS: 192, EBSS + Palm B: 126, Auto: 149, Auto + Palm B: 86) ** p<0.01, ***p<0.001. Scale bars: 10 µm.

**Figure 5.**
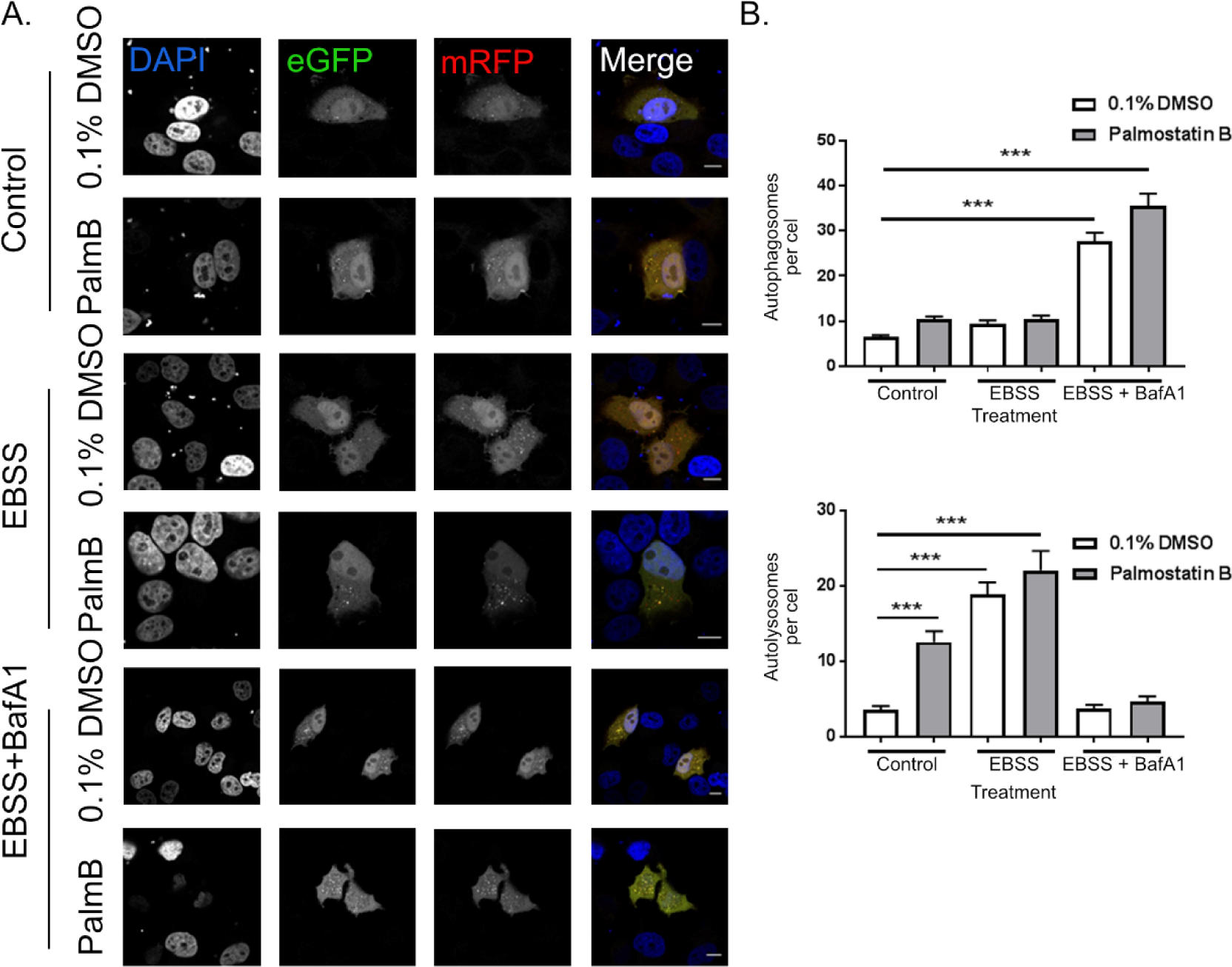
Palmostatin B alters autophagy in HeLa cells, assessed through the tandem LC3 reporter. HeLa cells were treated with 0.1% DMSO or 100μM Palmostatin (Palm. B) for 18h. Autophagy was induced by 4h EBSS treatment or induced and inhibited by 4h EBSS and 2h 500nM BafA1. **A.** Representative images of immunofluorescence conducted for eGFP-mRFP-LC3 after Palm. B and autophagy treatment. **B.** Quantification of the number of autophagosomes per cell (yellow puncta) and autolysosomes per cell (red puncta). Quantification was conducted on Image J software (mean + SEM from three independent experiments; Mann Whitney test; N (cells) Control: 123, Control + Palm. B: 64, EBSS: 71, EBSS + Palm B: 44, Auto: 85, Auto + Palm B: 64) ***p<0.001. Scale bars represent 10µm.

### S-acylated SQSTM1 is degraded by the lysosome

In contrast to LC3, SQSTM1 has not been shown to undergo any lipid modifications. However, it has been identified as potentially S-acylated in several S-acyl proteome studies^8,15^. Consequently, we sought to investigate if SQSTM1 undergoes S-acylation. HeLa cells were induced to undergo autophagy by serum deprivation in the presence or absence of BafA1 to prevent SQSTM1 degradation in the lysosome. S-acylation was detected using acyl-RAC. In the absence of autophagy or BafA1, a low level of S-acylation of SQSTM1 was detected (Figure 6). Serum deprivation led to a modest increase of SQSTM1 S-acylation with a concomitant decrease in total SQSTM1 protein suggesting that S-acylated SQSTM1 is directed to the lysosome where it is degraded. The addition of BafA1 significantly increased SQSTM1 S-acylation, further confirming that S-acylated SQSTM1 is directed to the lysosome but cannot be degraded when fusion between the lysosome and autophagosome are blocked with BafA1.

**Figure 6.**
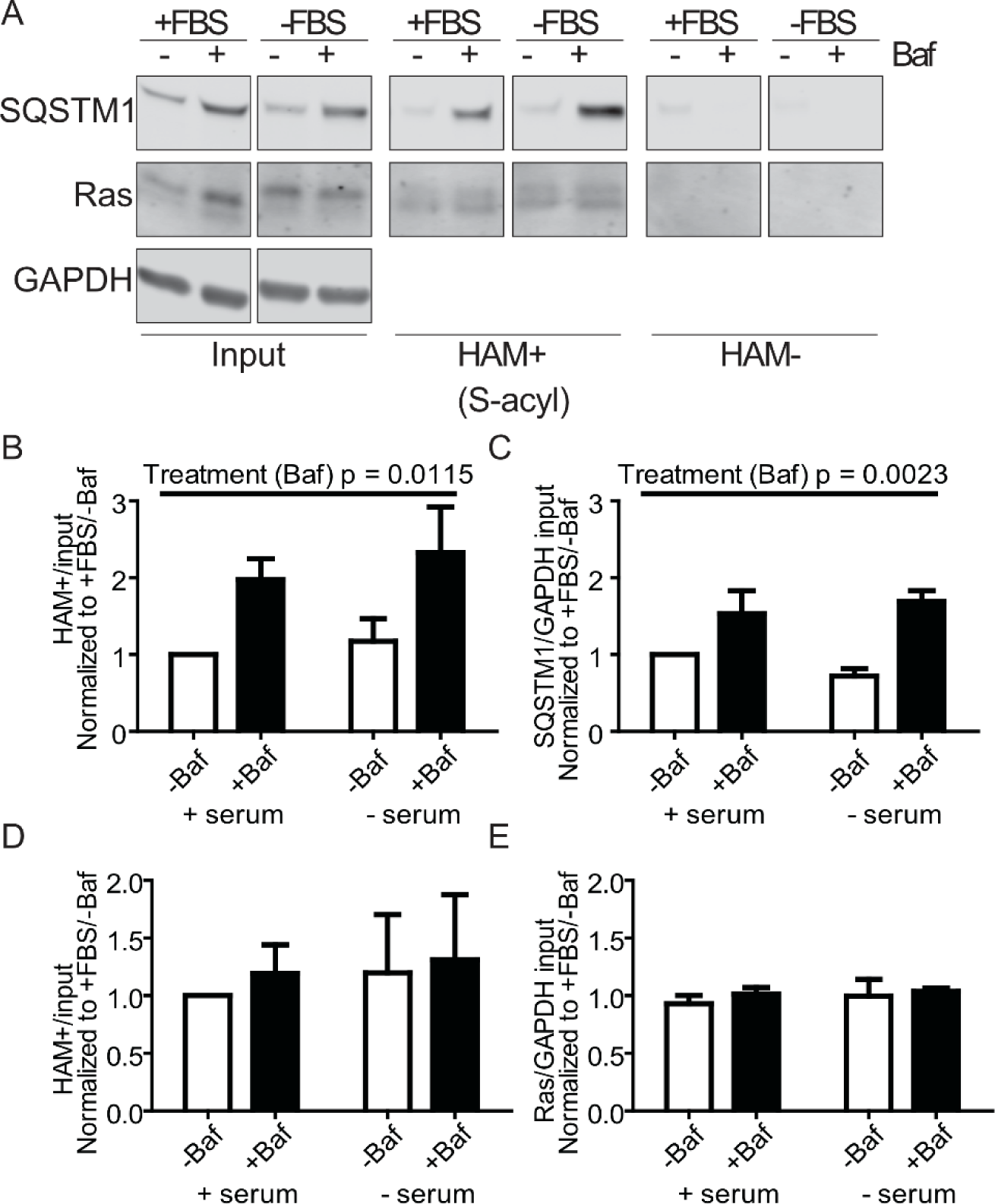
SQSTM1 is S-acylated and degraded by the lysosome. HeLa cells were induced to undergo autophagy through serum deprivation (-FBS) in the presence or absence of 400 nM BafA1. **A**. S-acylation of SQSTM1 and RAS was detected by the acyl-RAC assay. **B**. During autophagy SQSTM1 S-acylation increased while **C**. total SQSTM1 levels decreased (degraded by the lysosome). BafA1 significantly increased both **B**. SQSTM1 S-acylation and **C**. total protein. **D**. S-acylated RAS and **E**. protein levels were not significantly affected by either serum deprivation or BafA1 (2-way ANOVA, n=3, SEM).

### SQSTM1 S-acylation at the C289,290 di-cysteine motif is required for delivery to the lysosome

S-acylation of SQSTM1 is predicted (CSS-Palm 3.0 and 4.0)^28^ to occur at two di-cysteine motifs; C26,27 and C289,290. Consequently, cysteine to serine mutations were made at both sites. Serine is more structurally similar to cysteine than other amino acids. Only the mutations at C289,290 significantly decreased S-acylation of SQSTM1 (Figure 7A & 7B). However, SQSTM1 S-acylation was not completely blocked, suggesting additional sites of palmitoylation. Interestingly, palmitoylation did not increase in the presence of BafA1 in C289,290S suggesting that this may be the site that directs SQSTM1 to the phagophore (the sequestering compartment that is the precursor to the autophagosome) and, ultimately, the lysosome (Figure 7A and B). Similarly, C289,290S-SQSTM1-HA decreased colocalization of SQSTM1-HA with the lysosome marker CTSB-BFP in HeLa cells induced to undergo autophagy with rapamycin in the presence of BafA1 (Figure 7C).

**Figure 7.**
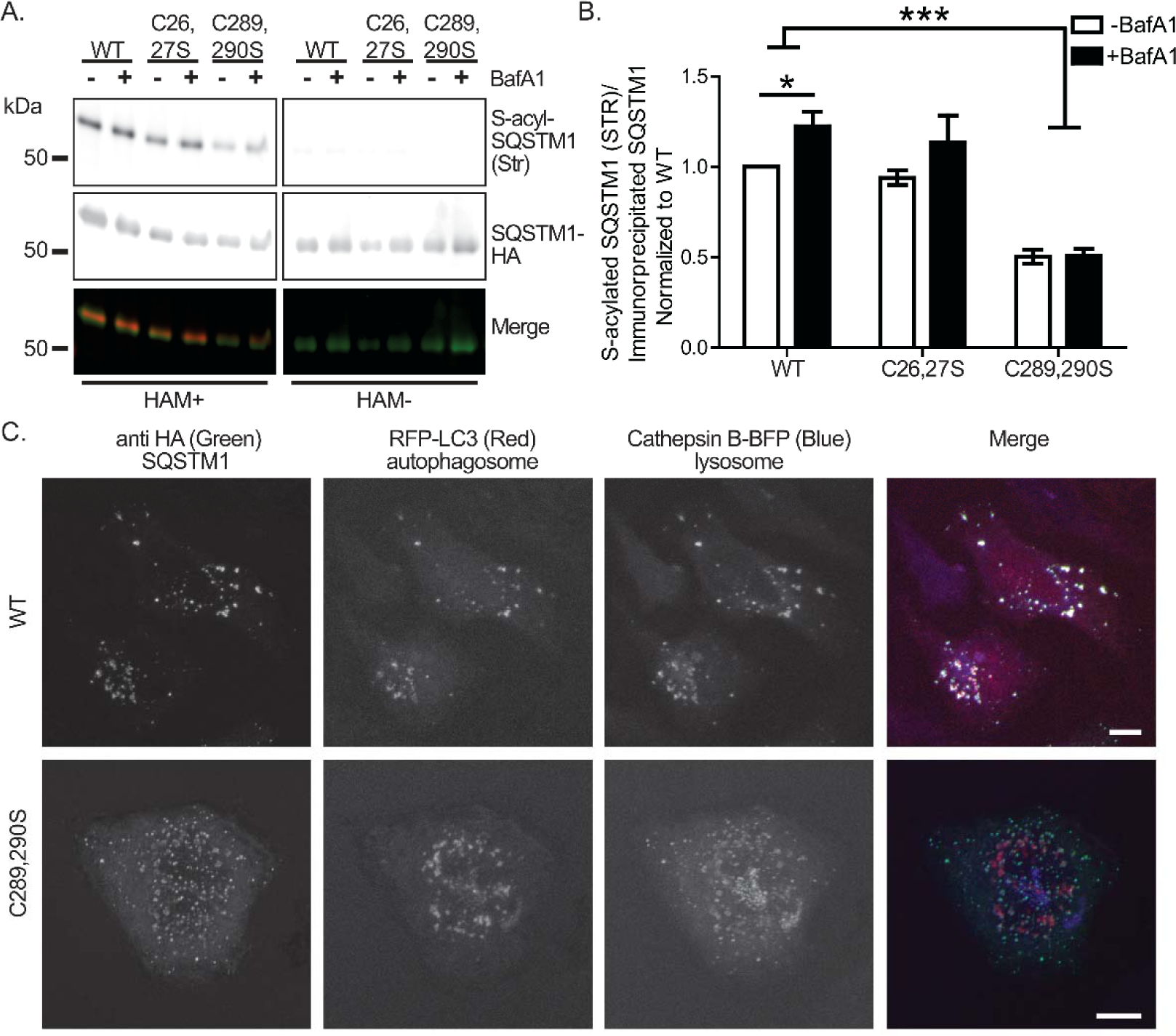
SQSTM1 is S-acylated at the di-cysteine C289, 290 motif which is required for localization to the lysosome. **A**. IP-ABE was performed from transfected HeLa cells expressing the indicated forms of SQSTM1-HA in the presence or absence of BafA1 and S-acylation was detected by streptavidin (STR). **B**. Only mutation of the C289, 290 di-cysteine motif significantly reduced S-acylation of SQSTM1-HA regardless of the addition of BafA1 (n = 4-7, *<0.05, *** <0.0001, 2-way ANOVA). **C**. HeLa cells expressing RFP-LC3 (autophagosome marker) and CTSB-BFP (lysosome marker) and the indicated SQSTM1-HA were treated with rapamycin to induce autophagy and BafA1 to block fusion and visualized by confocal microscopy.

### S-acylation of SQSTM1 is decreased in brains of HD patients

Several studies have indicated a cargo-loading defect in cells derived from HD models and patients^18,19,29^, leading to empty autophagosomes and a build-up of cytoplasmic debris thereby contributing to the pathogenesis of HD. As a crucial receptor for autophagy that targets aggregated proteins, mutant HTT (huntingtin; mHTT), and damaged mitochondria ^2,22^, we hypothesized that SQSTM1 S-acylation is reduced in HD which contributes to the empty autophagosome phenotype. As such, acyl-resin assisted capture was performed from donated HD patient brains to measure SQSTM1 S-acylation. S-acylation of SQSTM1 was significantly reduced in the cortex from HD patients compared to unaffected controls (Figure 8), despite a concomitant, but statistically insignificant, increase in total SQSTM1 levels. This result was also recapitulated in the brains from 8-month-old YAC128 HD mice by IP-ABE compared to wild-type (WT) control mice (Figure 9). Here, S-acylation was detected in the brains of 8 month old mice by ABE following immunoprecipitation of SQSTM1. Mice were either kept on a regular diet, or fasted overnight (24 h) to induce autophagy, as previously described^22^. Similar to HD patients, we detected a significant decrease in S-acylation of SQSTM1 in the YAC128 HD mouse brains (black bars – Fed) compared to their WT littermates (White Bars – Fed).

**Figure 8.**
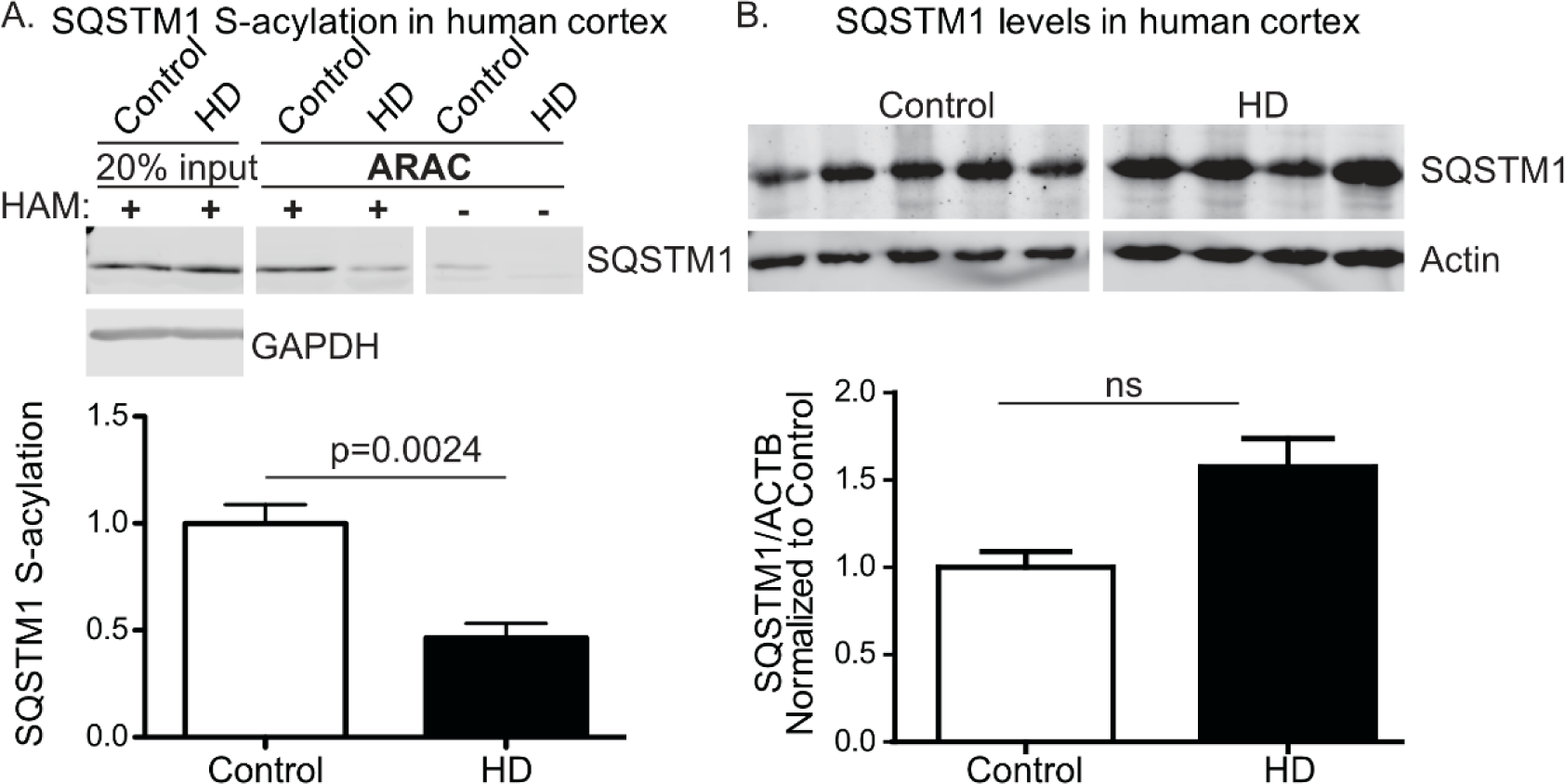
SQSTM1 palmitoylation is significantly reduced in HD patient brains. **A**. Palmitoylation of SQSTM1 was detected in patient brains by acyl-resin assisted capture. Tissues used were from patients of similar age at death, CAG size and post-mortem interval. Male and female samples were included in controls and HD samples. 2-tailed unpaired t-test, SEM (n=5,4 for Ctrl, HD). **B**. Total SQSTM1 levels were examined in HD patients. ns, not significant.

**Figure 9.**
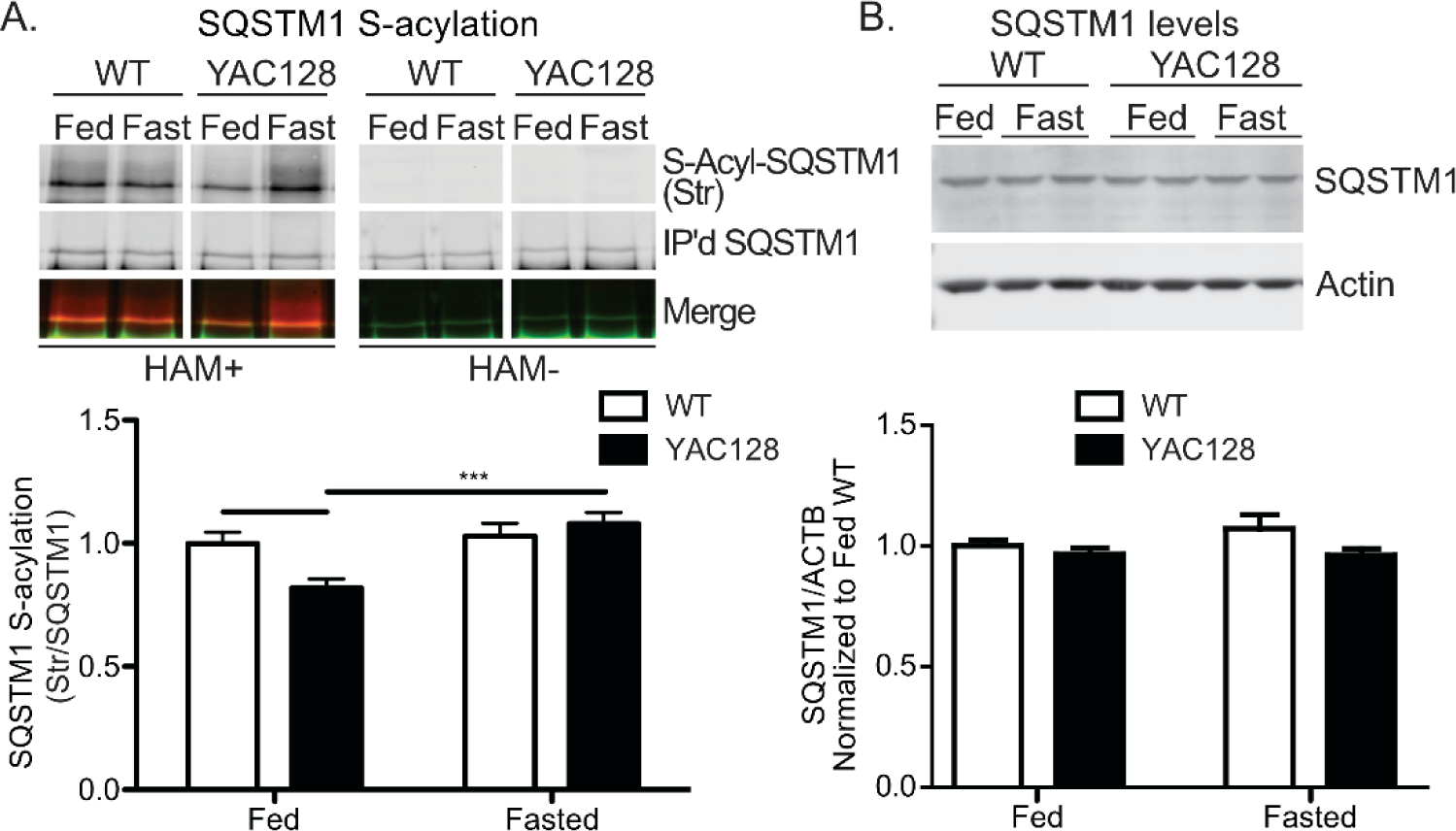
Significantly reduced SQSTM1 palmitoylation in the brains of HD mice is restored by fasting. Eight-month-old YAC128 mice were fasted for 24 h, as previously described. **A**. Palmitoylation of SQSTM1 was detected using IP-ABE (SQSTM1 is immunoprecipitated prior to ABE - top) and quantified (bottom). **B**) Total SQSTM1 levels were immunodetected (top) and quantified (bottom). Statistical significance was determined by 2-way ANOVA with Bonferroni’s post-hoc comparison; * p<0.05, *** p<0.0001. Data is represented as SEM.

### Fasting-induced autophagy rescues SQSTM1 acylation in the brains of the YAC128 HD mouse model

Previously, we demonstrated that autophagy could be induced through fasting in the YAC128 mouse^22^. As such, SQSTM1 palmitoylation was assessed in fed and 24-h fasted WT and YAC128 mice (Figure 9). Fasting rescued the decrease in SQSTM1 palmitoylation in YAC128 mouse brains to WT levels.

## Discussion

The significant enrichment of palmitoylated proteins in autophagy, as identified by our bioinformatics approach and increase of autophagy proteins in response to the depalmitoylation inhibitor palmostatin B, suggests that palmitoylation likely plays an important role in regulating autophagy. Palmitoylation likely regulates autophagy by acting as a mechanism for autophagy regulators to quickly localize to and from membranes during induction of autophagy. Recently, other regulators of autophagy have been shown to be palmitoylated, including, AKT, MTOR and LAMTOR1^17,30^. We predict that negative regulators of autophagy found at membrane interfaces, like MTOR, are palmitoylated when active and depalmitoylated when autophagy is activated. Conversely, we predict that autophagy regulators that need to quickly localize to membranes (i.e., LC3 lipidating enzymes, SQSTM1, etc.) are palmitoylated when autophagy is activated. Additionally, some proteins could also act as lipid sensors. For example, the LC3 lipidating enzymes have active cysteines that may be palmitoylated and inactive in the absence of autophagy. Fasting may promote their depalmitoylation and subsequent activation and lipidation of LC3.

Our data indicate two potential processes by which palmostatin B-mediated increases in palmitoylation may cause altered autophagy. Increased palmitoylation either causes an overall increase in autophagy or it causes a late-stage autophagy blockage. Although it is clear enhanced palmitoylation influences autophagy levels, we sought to examine the mechanism behind this. We found clear differences in autophagy proteins LC3 and SQSTM1 after upregulation of palmitoylation via palmostatin B treatment. Due to the important link between SQSTM1 and HTT, we chose to determine if SQSTM1 was in fact palmitoylated. Subsequently, we confirmed it is modified via palmitoylation (Figure 2) and showed that it is regulated by autophagy induction. Blocking the fusion between autophagosomes and lysosomes led to an increase in SQSTM1 levels and palmitoylation suggesting that lipidated SQSTM1 is directed to phagophores as is lipidated LC3. Although SQSTM1 has LC3-interacting motifs^31,32^, palmitoylation may provide the initial or additional signal that brings LC3 and SQSTM1 into proximity to interact. Palmitoylation may also provide an LC3-independent mechanism to direct SQSTM1 to the phagophore. Studying SQSTM1 palmitoylation is complicated by the fact that autophagosome-directed SQSTM1 is ultimately degraded by the lysosome. Therefore, it is difficult to detect SQSTM1 palmitoylation without blocking the fusion between the autophagosome and lysosome with compounds such as BafA1. In turn, inducing SQSTM1 palmitoylation presumably leads to degradation of SQSTM1, making it harder to detect.

Wildtype HTT is involved in regulating basal or bulk autophagy, while mHTT is associated with defects in autophagosomal and lysosomal fusion^2,29,32,33^. In combination with the known cargo-binding defect^19,32^, this is likely why inducing autophagy in HD is not efficient and potentially toxic, akin to adding water to a clogged sink. As such, this is why we determined that the best method to remove mHTT was to promote fasting-induced autophagy, an HTT-independent pathway^22^. This may partly explain why mHTT is lowered in the brains of YAC128 mice when autophagy is induced through a week-long intermittent fasting paradigm. Similarly, herein, a short fasting time increased the palmitoylation of mHTT in the brains of YAC128 mice. Therefore, fasting-induced autophagy may also correct the cargo loading defect seen in HD by increasing SQSTM1 palmitoylation and localization to the autophagosome and lysosome for degradation.

Similar to SQSTM1, we have previously shown that mutant HTT palmitoylation is lower in several models of HD, including patient-derived lymphoblastoid cells and stem cells^25,34^. Blocking depalmitoylation with the broad inhibitor palmostatin B decreased insoluble mutant HTT and increased HTT palmitoylation with a concomitant increase in cell survival in primary neuronal cells derived from the YAC128 HD mouse model^25^. An independent study verified the protective effects of blocking depalmitoylation^35^. Although the latter study did not directly measure palmitoylation, these studies^25,35^ combined suggest that promoting palmitoylation in HD is protective. The protective effect may also be due to increased SQSTM1 palmitoylation that directs mutant HTT to the autophagosome for degradation. This would be an interesting area of future studies.

Ultimately, the data presented suggest that approaches that promote SQSTM1 palmitoylation may be beneficial in HD. Of note, many mutations in SQSTM1 are associated with ALS, frontotemporal dementia, and Charcot-Marie Tooth disease^36^, and SQSTM1 deposits are also hallmark features of ALS, Charcot-Marie Tooth disease, and multisystem proteinopathy^37,38^. Therefore, palmitoylation of SQSTM1 may be a potential marker for disease, and modulating SQSTM1 palmitoylation may be an attractive therapeutic approach in multiple disorders. It will be important to investigate if SQSTM1 palmitoylation is altered in these diseases and how it is regulated.

## ACKNOWLEDGEMENTS

The authors would like to thank members of the Hayden lab for helpful discussions and input. In particular, Mahsa Amir, Mark Wang, and Yun Ko for organizing and maintaining mouse colonies and Dr. Amber Southwell for helpful comments for mouse studies. Thank you to Dr. Klionsky for your valuable feedback on the manuscript. Finally, we would like to thank all the patients and their families who donated tissues to help move HD research forward. This work could not have been done without them. MC and MD were supported by a grant from The Royal Society (RG150409). DDOM, FA, AK, and AD were supported by a Natural Sciences and Engineering Research Council (NSERC) Discovery Grant (RGPIN-2019-04617)

